# A process-based model simulating the life cycle of *Culex pipiens* s.s./*Cx. torrentium* in Germany

**DOI:** 10.1101/2024.11.20.624534

**Authors:** L. Rauhöft, P. Duve, S.M. Martins Afonso, H. Jöst, T. Şuleşco, F.G. Sauer, R. Lühken

## Abstract

Mosquitoes are well known for their ability to transmit pathogens, including various arthropod-borne viruses (arboviruses) of veterinary and medical interest. The threat of (re-)emerging arboviruses in Europe is increasing due to globalization and climate warming. This also applies to temperate regions, where the transmission of viruses is becoming possible due to an increase of the ambient temperature, shortening of the extrinsic incubation period. *Culex pipiens* s.s./*Cx. torrentium* are the primary vectors of Usutu virus and West Nile virus in Europe and are found in and around human settlements. The prediction of spatial-temporal abundance allows for the early assessment of arbovirus transmission risk and the planning of effective intervention methods, such as vector control. Therefore, a process-based model was developed to predict the spatial-temporal occurrence of *Cx. pipiens* s.s./*Cx. torrentium* in Germany with a particular focus on depicting realistic overwintering behaviour, e.g. diapause induced through photoperiod and temperature in the larval stage. The model output is driven by local temperature and rainfall data. Evaluated with field data from 116 sampling sites in Germany, the model accurately identified the peak in abundance, with a mean absolute off-set of 0 days between the simulated and observed peak, and offsets of 8 and 1 days for the start and end of the mosquito season, respectively. A significant linear relationship between simulated and observed mosquito abundance was found for 78.45% of the sampling sites, with an overall significant linear relationship across all sites (Estimate = 0.16, Standard Error = 0.003, *t*-value = 48.54, degrees of freedom = 2724, *p*-value < 0.0001, marginal R^2^= 0.47). This model offers a robust framework for the prediction of the mosquito population dynamics of *Cx. pipiens* s.s./*Cx. torrentium* under current and future climate scenarios, thereby supporting vector surveillance and control strategies across Europe.

**Graphical Abstract:** 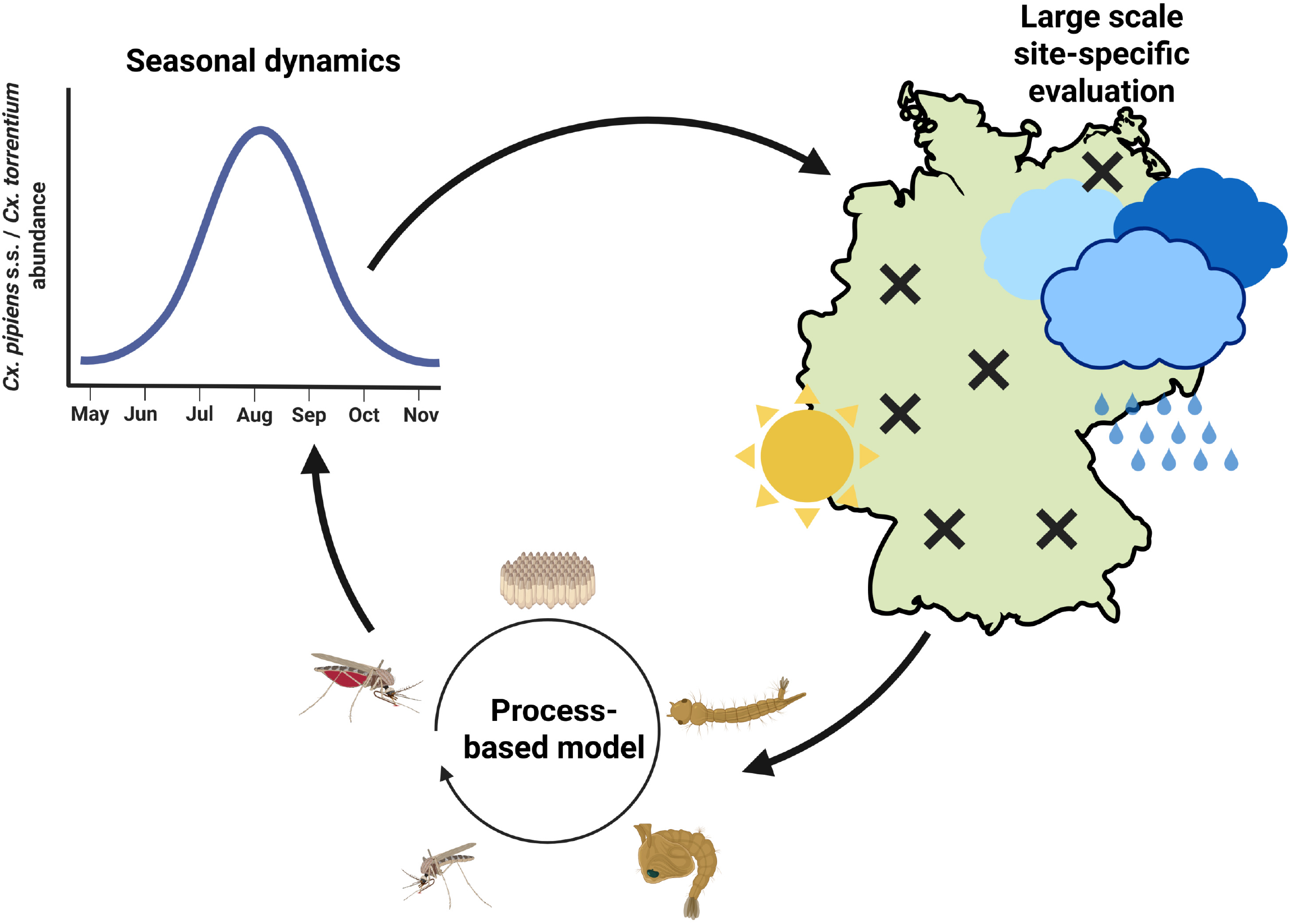

**Highlights:**

- Accurate depiction of the phenology of *Culex pipiens* s.s./ *Cx. torrentium*, including the time point of the start, end and peak of the mosquito season.
- Large-scale and site-specific validation using nation-wide mosquito abundance data from 116 sampling sites from five collection years, indicated a significant linear relationship between field data and model output for 78.45% of the sampling sites.
- First biologically accurate simulation of the overwintering behaviour of *Culex pipiens* s.s./ *Cx. torrentium*.
- Europe-wide raster map showing the maximum number of consecutive generations per season.

## 1. Introduction

Mosquitoes of the species *Culex pipiens* s.s./*Cx. torrentium* are of major importance as vectors of medical and veterinary relevant arboviruses (Brugman et al., 2018). They have a high vector competence for Usutu virus (USUV) and West Nile Virus (WNV) (Jansen et al., 2019; Fros et al., 2015). *Cx. pipiens* s.s./*Cx. torrentium* are widely distributed and have broad host-feeding patterns, including birds, non-human mammals and humans, making them suitable bridge vectors (Wehmeyer et al., 2024). Therefore, these species are a main target for surveillance and risk assessment efforts (Ros et al., 2014; Rudolf et al., 2013).

Mosquito collections provide the basis to identify the spatial-temporal risk of mosquito-borne pathogen transmission. The collection, mostly using CO_2_-baited traps, and the subsequent taxa identification and screening for arboviruses in the laboratory is time-consuming and costly. Process-based models can offer a valuable alternative to predict mosquito populations and transmission dynamics, using mathematical equations to simulate natural phenomena, as applied here by the life cycle of *Cx. pipiens* s.s./*Cx. torrenium*. Driven by environmental factors, especially temperature and rainfall, these models enable the estimation of the spatial-temporal mosquito abundance and at the same time also allow the simulation of changing conditions, e.g. control measurements (Cailly et al., 2012). Several process-based models for mosquitoes have been published for Europe focussing on the Mediterranean region, i.e. Southern France or Italy (Cailly et al., 2012; Ezanno et al., 2015; Tran et al., 2013; Marini et al., 2016). These models showed a high performance in the prediction of the temporal mosquito abundance, but were generally only validated with field data from few sampling sites or relatively small areas. Thus, a process-based model for *Cx. pipiens* s.s./*Cx. torrentium* as one of the most important vectors in Central Europe and a large-scale, site-specific evaluation is missing. In this work, we developed and validated a process-based model for *Cx. pipiens* s.s./*Cx. torrentium* in Germany. To achieve this, we adjusted the generic model for *Ae. albopictus* originally proposed by Cailly et al. (2012) and later extended to further species by Ezanno et al. (2015) (Figure 1). Firstly, temperature-dependent transition functions during the mosquito life cycle were adapted to broaden the temperature range in which the model is applicable, thereby expanding its geographic range of usability including Central Europe. Secondly, in contrast to previous studies, our model no longer relies on a predefined favourable and unfavourable season to determine the start and the end of the overwintering period (Tran et al., 2013; Ezanno et al., 2015). This predefined binary variable, based on expert knowledge, triggers one of two sets of equations where mosquitoes are either active or in overwintering. The diapause duration can substantially influence the transmission risk of pathogens, since this influences the duration of the mosquito biting season. Furthermore, models which do not accurately predict the biting season can only give limited predictions of disease risk (Ewing et al., 2021). This system therefore was replaced by new compartments for the overwintering (diapausing) stages. Since the diapause of female *Cx. pipiens* s.s./*Cx. torrentium* is triggered during the larval stage (Robich and Denlinger, 2005), we extended the model by sex-differentiated compartments, i.e. male and female eggs, larvae and pupae. The latter females further split into a diapause-conditioned pupal stage, which then develops into an emerging diapausing adult stage that mates before overwintering. In future, the sex-differentiated compartments also allow the analysis of different mosquito control measures such as the sterile insect technique or Wolbachia-based mosquito suppression or reduction of vector competence (Anguelov et al., 2017; Bourtzis et al., 2014).

**Figure 1:**
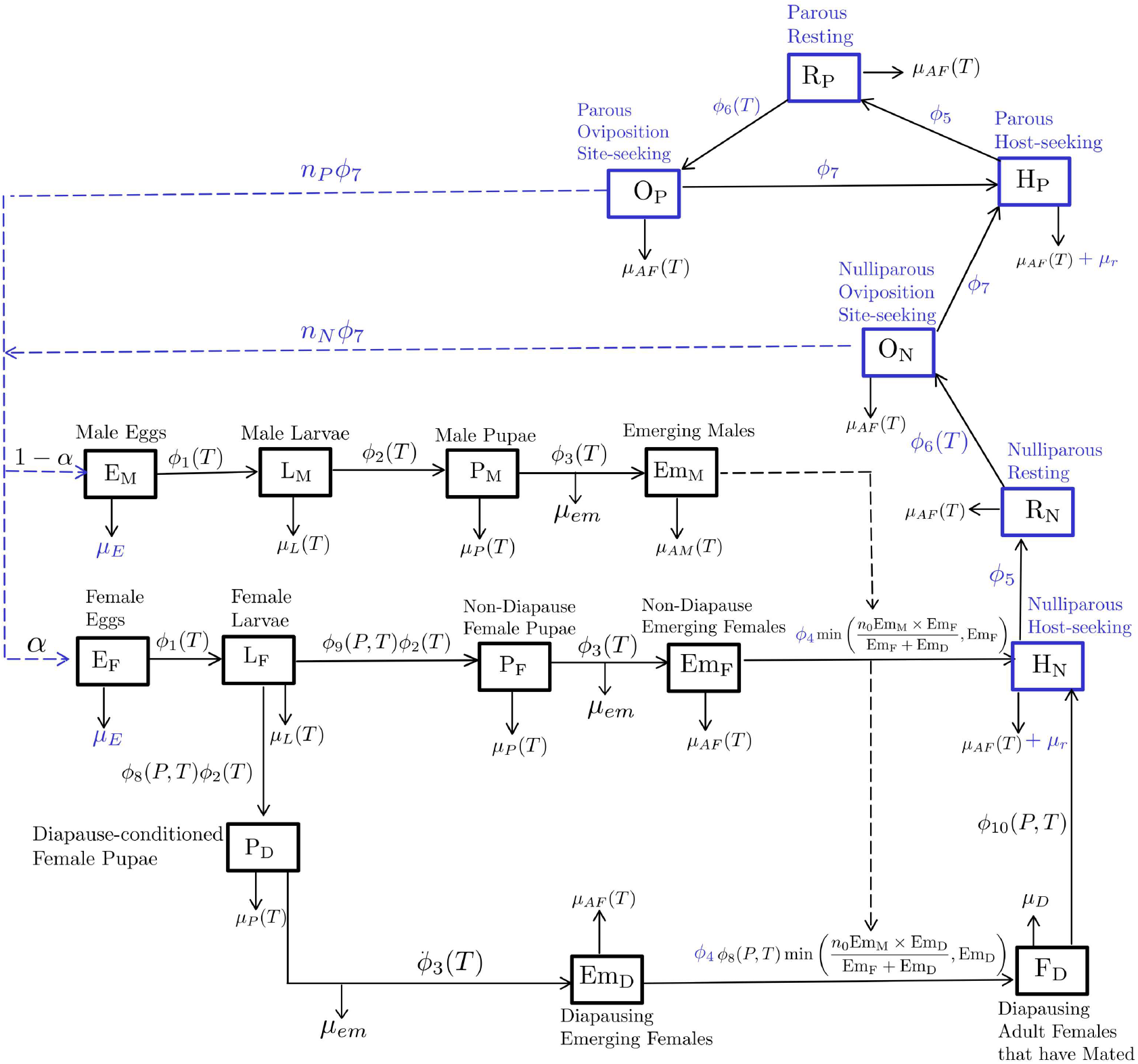
Flow chart diagram for the process-based model for the mosquito life cycle of *Cx. pipiens*, s.s./,*Cx. torrentium*). Boxes represent compartments or different life stages. Solid lines represent the development from one stage to another and dashed lines represent the interactions between the different compartments. Previously published compartments, transition and mortality rates are highlighted in blue.

The new process-based model was validated with field-data for Germany, including the start, end and peak of annual mosquito activity. Furthermore, the prediction accuracy of the model was compared to that of a temperature-dependent correlative model. The variable importance for the overall mosquito dynamic and the population peak was analysed using a sensitivity analysis.

## 2. Methods

### 2.1. Mathematical model for the life cycle of *Cx. pipiens* s.s./*Cx. torrentium*

The model consists of the different compartments for mosquito abundance of aquatic stages: male and female eggs (*E*_*m*_, *E*_*f*_), larvae (*L*_*M*_, *L*_*F*_), pupae (*P*_*M*_, *P*_*F*_), and diapause-conditioned pupae (*P*_*D*_), and adult stages: emerging males and females (*Em*_*M*_, *Em*_*F*_), nulliparous host-seeking females (*H*_*N*_), nulliparous resting females (*R*_*N*_), nulliparous oviposition site-seeking females (*O*_*N*_), parous host-seeking females (*H*_*P*_), parous resting females (*R*_*P*_) and parous oviposition site-seeking females (*O*_*P*_) and diapausing stages: emerging adult females (*Em*_*D*_) and overwintering adult females (*F*_*D*_)(Equation 7). Transition between the aquatic stages is temperature-dependent (*ϕ*_1_(*T*), *ϕ*_2_(*T*), *ϕ*_3_(*T*); Equation 1). Previous laboratory and semi-field studies suggest that diapause is induced at photoperiods lower than 15 hours and temperatures under 20^°^C (Spielman and Wong, 1973; Field et al., 2022). This is represented in our temperature- and photoperiod-dependent function *ϕ*_8_(*P, T*) inducing diapause (Figure 2 B; Equation 5). The rate of diapause termination is also temperature- and photoperiod-dependent (*ϕ*_10_(*P, T*); Equation 6; Nelms et al. (2013)). Additionally, temperature-dependent mortality rates were utilized for larvae and pupae (*µ*_*L*_(*T*), *µ*_*P*_ (*T*); Equation 2; Figure 3). Rainfall increases the number of available breeding sites. The competition for resources within the breeding site is taken into account by the density-dependent larval mortality of 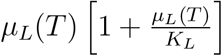(Ezanno et al., 2015; Legros et al., 2009; Lutambi et al., 2013). The pupal density dependence was suggested to be combined in an exponential decay function using the mortality at emergence (*µ*_*em*_) as scaling factor. Firstly, we propose that the mortality at emergence (*µ*_*em*_) is not a matter of resources since pupae do not take up any nutrients. Thereby the mortality at emergence can only be a stochastic event or a matter of actual space restriction by density. We thereby replaced the scaling factor with *β* and strongly reduced it compared *µ*_*em*_. The mortality at emergence was kept outside of the exponential decay function. A proportion *α* of mosquito eggs are female while a proportion of (1 − *α*) are male. Based on the assumption that all female mosquitoes mate before seeking for a host, emerging females (*Em*_*F*_) and emerging diapausing females (*Em*_*D*_) mate before the females progress either to the host-seeking stage (*H*_*N*_) or the over-wintering stage (*F*_*D*_). As also implemented by Ezanno et al. (2015), it is assumed that females mate only once. In contrast, males can mate several times, i.e. Asman (1975) showed that under laboratory conditions male *Cx*.*tarsalis* mosquitoes can inseminate up to 8 females over 21 days. Thereby, the number of females which get fertilized by males is highly dependent on the population density of sexually active male mosquitoes (Anguelov et al., 2017). When sexually active males are scarce in the environment, the number of inseminated females that progress to the host-seeking stage is proportional to the number of available males multiplied by the number of females each male can inseminate. This is denoted as *n*_0_*Em*_*M*_, where *n*_0_ is the number of females each male inseminates. After mating, female mosquitoes progress to the host-seeking stage at a rate of *ϕ*_4_. Blood feeding induces the resting stage at a rate of *ϕ*_5_ before females start seeking an oviposition site to lay eggs at a rate of *ϕ*_6_(*T*) (Equation 4). Female mosquitoes become parous after depositing eggs at a rate of *ϕ*_7_. The cycle repeats until the mosquitoes die and are removed from the population. During the diapausing stage, the overwintering females die with a rate of *µ*_*D*_. Female adult mosquitoes in general die at a rate of *µ*_*AF*_ (*T*) (Equation 2), while for host-seeking females *H*_*N*_ and *H*_*P*_ an additional host seeking mortality *µ*_*r*_ is added. Adult males are shown to have shorter longevity especially during the highest and lowest temperatures (Andreadis et al., 2014). Therefore, an additional mortality rate for males *µ*_*AM*_ (*T*) (Equation 3) was generated based on the general adult mortality rate but more constrained in the temperature range (Figure 3).

### 2.2. Validation

Data for the validation of the process-based model was collected with CO_2_ -baited mosquito traps on weekly, bi-weekly or daily basis from April to October in the years 2016-2017 and 2021-2023 including many sites run by voluntary helpers (Figure 9). All mosquitoes were identified by morphology using the taxonomic key by Becker et al. (2020). Additionally, the blood-fed status of the caught mosquitoes was assessed using the Sella score (Detinova, 1962), categorizing mosquitoes as unfed (Sella score 1) or as blood-fed from freshly engorged to gravid (Sella score 2-7). Only female mosquitoes with a Sella score of one were used for validation, since these are the specimens considered being host-seeking. Only sampling sites with more than six sampling dates per year, a total number of more than 30 captured *Cx. pipiens* s.s./*Cx. torrentium* females per year and more than 10 captured during the seasonal peak were included in the analysis, as this threshold was considered the minimum necessary to catch a representative seasonal dynamic. The final validation data set consisted of 116 sampling sites (Supplementary Table S1) with 77,535 *Cx. pipiens* s.s./*Cx. torrentium* females (mean per day = 24.98, 95% confidence interval (CI95) = 20.19 - 29.76).

**Figure 2:**
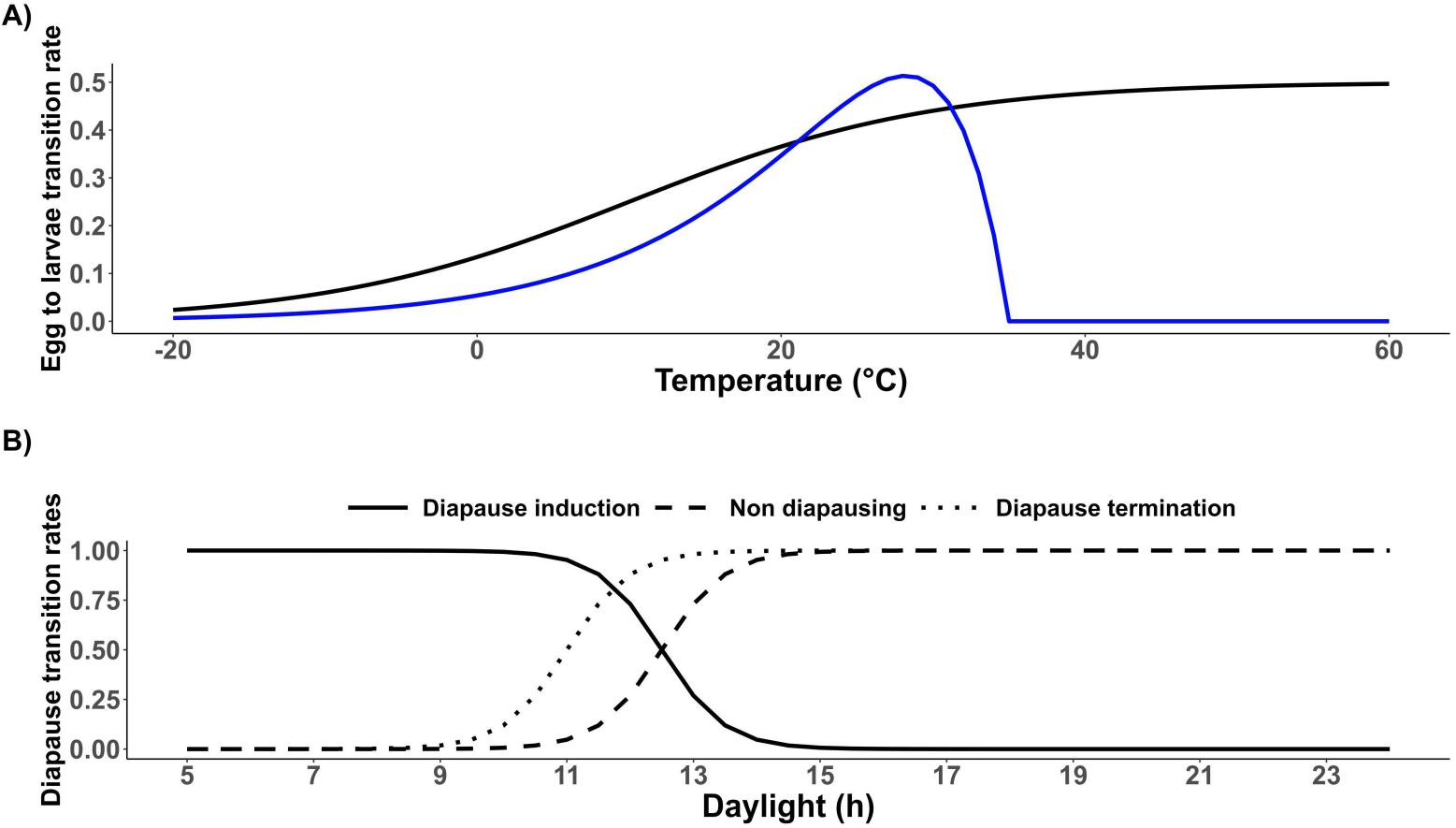
A) Comparison of the former (blue) (Ezanno et al., 2015) and newly developed (black) temperature-dependent transition rates and B) newly developed photoperiod-dependent transition rates.

**Figure 3:**
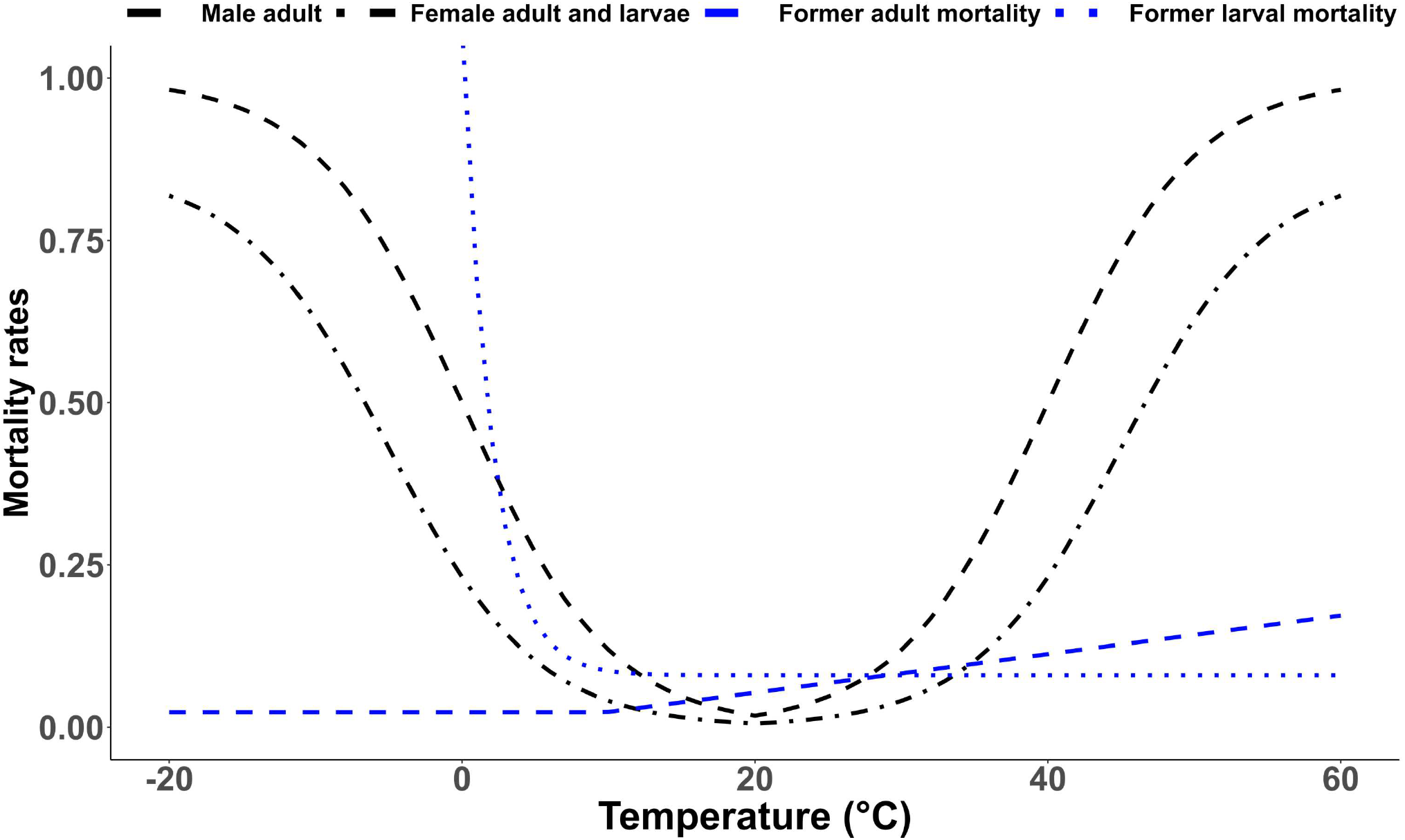
Comparison of the former (blue) (Ezanno et al., 2015) and newly developed (black) temperature-dependent mortality rates.

Mean daily temperature and daily rainfall data (European re-analysis and observations for monitoring, E-OBS, v29.0e) were obtained from the ECA&D project (http://www.ecad.eu, Cornes et al. 2018) and extracted for each sampling site for 2015 - 2023. For each sampling site and sampling date, the sum of predicted nulliparous and parous host-seeking females was compared with data from the field. Therefore, smoothed rolling means were calculated for both, the captured mosquitoes and the model output with a rolling window length of four days, as both can exhibit strong short-term fluctuations. For each sampling site, linear models (LMs) were calculated between model prediction and field capture, after log transforming the field captures and adding 0.01 due to the high numbers of zeroes in the data. Mosquito phenology is strongly influenced by temperature. Therefore, as suggested by Ezanno et al. (2015) LMs were also calculated between daily mean temperature and observed field captures for each sampling site to analyse how well temperature data alone explains the observed mosquito abundance. In this case also 0.01 was added to the log transformed field data. A Student’s t-test was used to determine if the correlation between the model’s predictions and actual mosquito counts from field data was significantly different from the correlation between temperature alone and field data. For each sampling site and sampling year, the offset between the abundance peaks was calculated as the difference between field-observed dates and model-predicted dates. The same was done for the begin and end of the season, defined as the time point where the number of mosquito specimens firstly and lastly exceeded 1% of the corresponding seasonal peak for each sampling site and year. Using the offsets, mean, median and 95% confidence intervals were calculated for field and model-predicted data. To assess the accuracy of the predicted seasonal peak compared to the observed field data a linear regression was performed. This analysis was done using the raw data. The same was done for the begin and end of the season, defined as the time point where the number of mosquito specimens firstly and lastly exceeded 1% of the corresponding seasonal peak for each sampling site and year. Validation for all available data was conducted with a linear mixed model (LMM), analyzing the number of predicted host-seeking female mosquitoes from the model and the log-transformed number of field-collected female mosquitoes with a Sella score of 1, after adding 0.01 due to the high number of zeroes in the data. Hereby, the sampling site was used as random factor. Again, we used an additional model to examine whether the process-based model provided any advantage over simply predicting mosquito phenology based solely on temperature data, which in our case was an LMM.

### 2.3. Consecutive generations per season

Considering the importance of the length of the mosquito biting season, all transition rates within the process-based model were extracted and used to calculate the developmental time to reach the next life stage. A seven-day average temperature at the beginning of each life stage was used for this calculation. In this manner we calculated a maximum possible number of consecutive generations per season which in turn can be used as a proxy for the length of the mosquito season. This model calculation was run over the entire Europe-wide E-OBS raster for the years 2021 to 2024. Due to known issues in the E-OBS data set near the edges (Copernicus Climate Change Service, 2024), the map was visually inspected, and regions with nonsensical high or low values were manually removed before calculating the area weighted minimum, mean and maximum number of consecutive generations for entire Europe. Additionally, we used a buffer zone of 50 km within the national borders before calculating the area weighted minimum, mean and maximum number of consecutive generations for Sweden, Germany and Spain.

### 2.4. Sensitivity analysis

We conducted a sensitivity analysis to gain insights into factors that influence the peak of the population of host-seeking females (sum of nulliparous and parous females) and the general phenology of the population of host-seeking females throughout the simulation period. Thereby, parameters are resampled using the Latin Hypercube Sampling method, which is an efficient stratified Monte Carlo sampling that allows for simultaneous sampling of the multidimensional parameter space (Blower and Dowlatabadi, 1994; Hoare et al., 2008). The simulation was carried out for a period of 1 year, with 1000 simulations per run. Partial rank correlation coefficients (PRCCs) are computed for each selected input parameter and the output variables (peak and phenology of host-seeking females) for the entire simulation period.

### 2.5. Software

All analysis was performed in R (Version: 4.4.0) using the R-Studio IDE (Version:2024.12.1) (R Core Team, 2024). Additionally, functions from the following packages were used for data preparation, visualization and analysis: lme4 (Bates et al., 2015), lmerTest (Kuznetsova et al., 2017), performance (L decke et al., 2021), dplyr (Wickham et al., 2023a), tidyr (Wickham et al., 2024), ggplot2 (Wickham, 2016), scales (Wickham et al., 2023b), rstatix (Kassambara, 2023), stringr (Wickham, 2023), deSolve (Soetaert et al., 2010), sensitivity (Pujol et al., 2008), lhs (Carnell, 2012), knitr (Xie, 2013), plyr (Wickham, 2011), ggpubr (Kogure et al., 2019), zoo (Zeileis and Grothendieck, 2005), raster (Hijmans, 2023), geosphere (Hijmans, 2024), lubridate (Grolemund and Wickham, 2011) and xtable (Dahl et al., 2019).

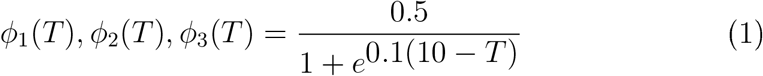

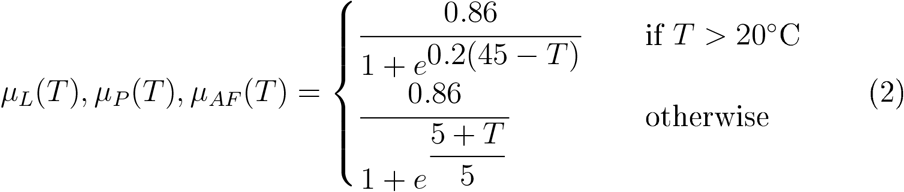

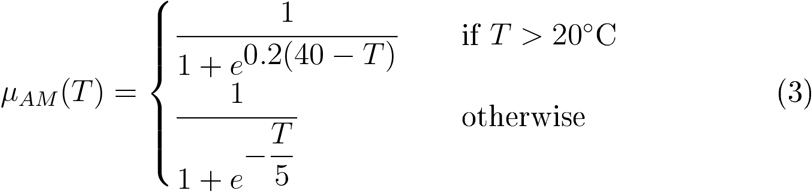

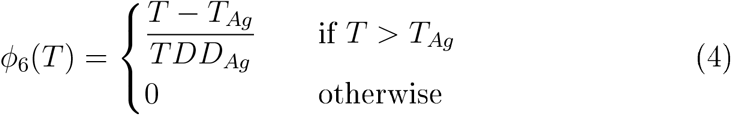

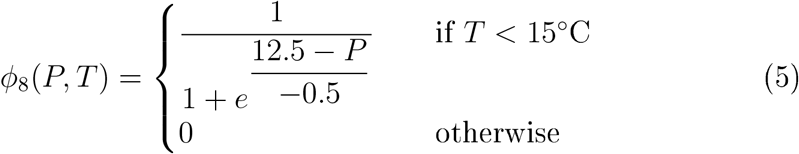

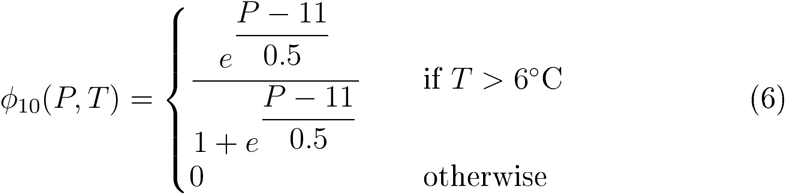

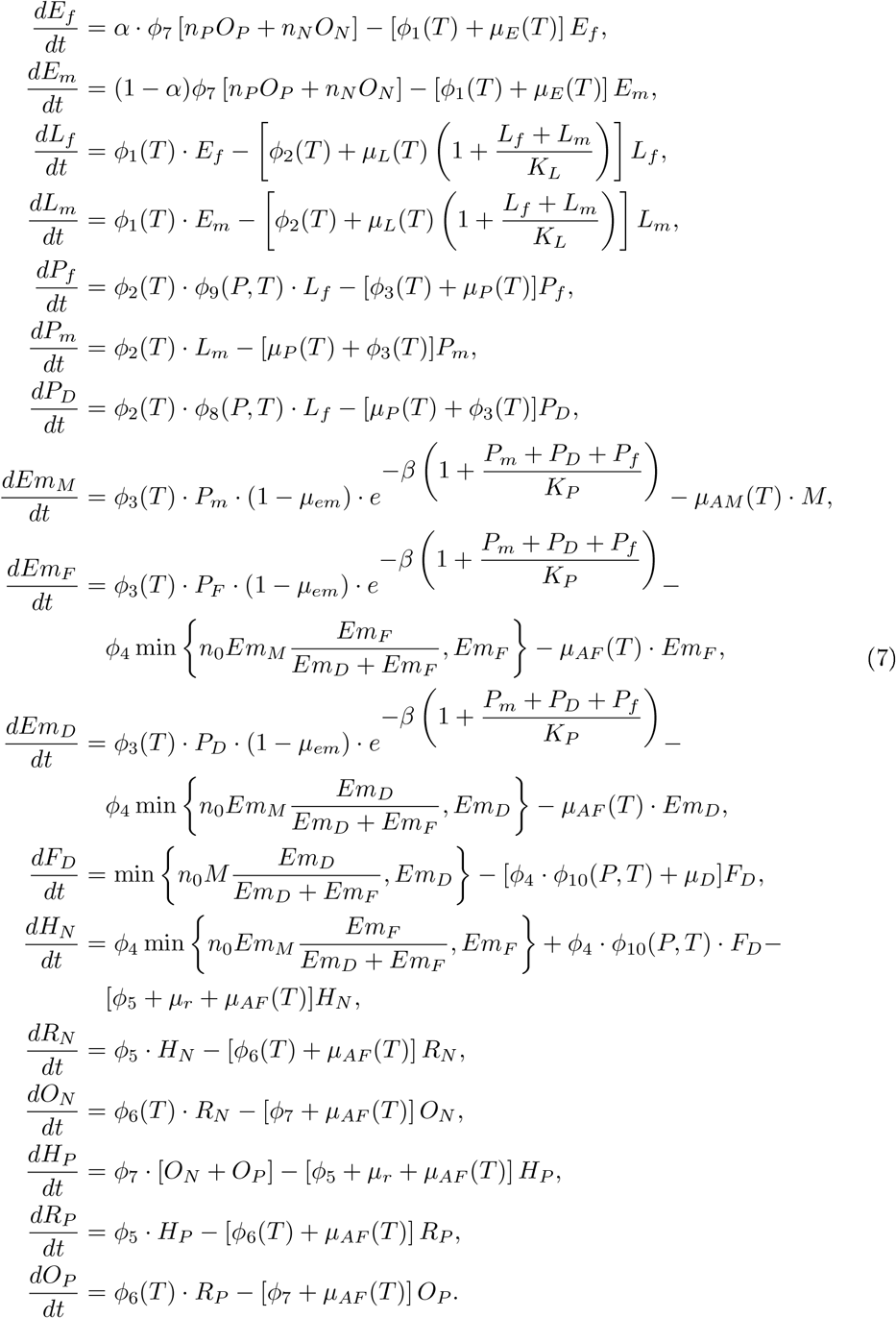

**Table 1:** Table of all used parameter, their values, formulas and origin

| Parameter | Definition | Value | Reference |
| --- | --- | --- | --- |
| $\phi_1(T)$ | Egg development rate | Eq. 1 | adopted from unpublished data |
| $\phi_2(T)$ | Larval development rate | Eq. 1 | adopted from unpublished data |
| $\phi_3(T)$ | Pupal development rate | Eq. 1 | adopted from unpublished data |
| $\phi_4$ | Host-seeking rate | 0.4 | Tran et al. (2013) |
| $\phi_5$ | Resting rate | 0.2 | Tran et al. (2013) |
| $\phi_6(T)$ | Oviposition site-seeking rate | Eq. 4 | Tran et al. (2013); Ezanno et al. (2015) |
| $\phi_7$ | Egg-laying rate | 0.2 | Tran et al. (2013) |
| $\phi_8(P, T)$ | Diapause entering | Eq. 5 | Spielman and Wong (1973); Field et al. (2022) |
| $\phi_9(P, T)$ | Non-diapause maintenance rate | $1 - \phi_8(P, T)$ | Spielman and Wong (1973); Field et al. (2022) |
| $\phi_{10}(P, T)$ | Diapause exiting rate | Eq. 6 | Nelms et al. (2013) |
| $\mu_E$ | Egg mortality rate | 0.0262 | Ezanno et al. (2015) |
| $\mu_L(T)$ | Larval mortality rate | Eq. 2 | adopted from unpublished data |
| $\mu_P(T)$ | Pupal mortality rate | Eq. 2 | adopted from unpublished data |
| $\mu_{AF}(T)$ | Mortality rate of female adult mosquito | Eq. 2 | adopted from unpublished data |
| $\mu_r(T)$ | Host-seeking mortality | 0.08 | Ezanno et al. (2015) |
| $\mu_{AM}(T)$ | Mortality rate of male adult mosquito | Eq. 3 | adopted from unpublished data |
| $\mu_D(T)$ | Mortality rate of diapausing female adult mosquito | 0.02 | adopted from unpublished data |
| $\alpha$ | Sex ratio at the emergence stage | 0.5 | Altizer et al. (2013); Ezanno et al. (2015) |
| $\beta$ | Strength of pupal density dependence | 0.003 | adopted from data |
| $T_E$ | Minimal temperature for egg maturation (°C) | 9.8 | Ahumada et al. (2004); Balenghien et al. (2008); Ezanno et al. (2015) |
| $TDD_{Ag}$ | Degree-days for egg maturation (°C) | 64.4 | Ahumada et al. (2004); Balenghien et al. (2008); Ezanno et al. (2015) |
| $K_l$ | Larval environmental carrying capacity | $8 \times 10^8$ | Ahumada et al. (2004); Balenghien et al. (2008) |
| $K_p$ | Pupal environmental carrying capacity | $10^7$ | Ahumada et al. (2004); Balenghien et al. (2008) |

## 3. Results

### 3.1. Validation

The process-based model predicted a seasonal dynamic for every life stage for each sampling site (Figure 4 B & C). With an average R^2^ -value of 0.53 (CI95 = 047 - 0.59), site specific LMs demonstrated a significant positive linear relationship between the field data and model output for 91 sampling sites (Supplementary Table S1). The remaining 25 sites, showed no significant correlation. In contrast, site specific LMs only correlating temperature with mosquito field data had an average R^2^-value of 0.33 (CI95 = 0.27 - 0.38). The process-based model explains significantly more variation in the eld data compared to the daily average temperature alone (*t*_(243)_ = 5.41, *p*-value (*p*) < 0.0001). The LMM over all sites, relating field captures to model predictions was significant (Estimate (E) = 0.16, Standard error (SE) = 0.003, *t*-value (*t*) = 48.54, degrees of freedom (df) = 2724, *p* < 0.0001) and explained 46.72% for the fixed effect only (marginal (marg.) R^2^= 0.47 and 55.77%, accounting for random variation among sampling sites (conditional (cond.) R^2^= 0.56). Again, the correlation between temperature and the eld data was significant (E = 1.66, SE = 0.05, *t* = 35.86, df = 2723, *p* < 0.0001, but explained less variation: 32.75% for the fixed effect (marg. R^2^ = 0.33) and 42.95% accounting for random variation among sampling sites (cond. R^2^= 0.43).

**Figure 4:**
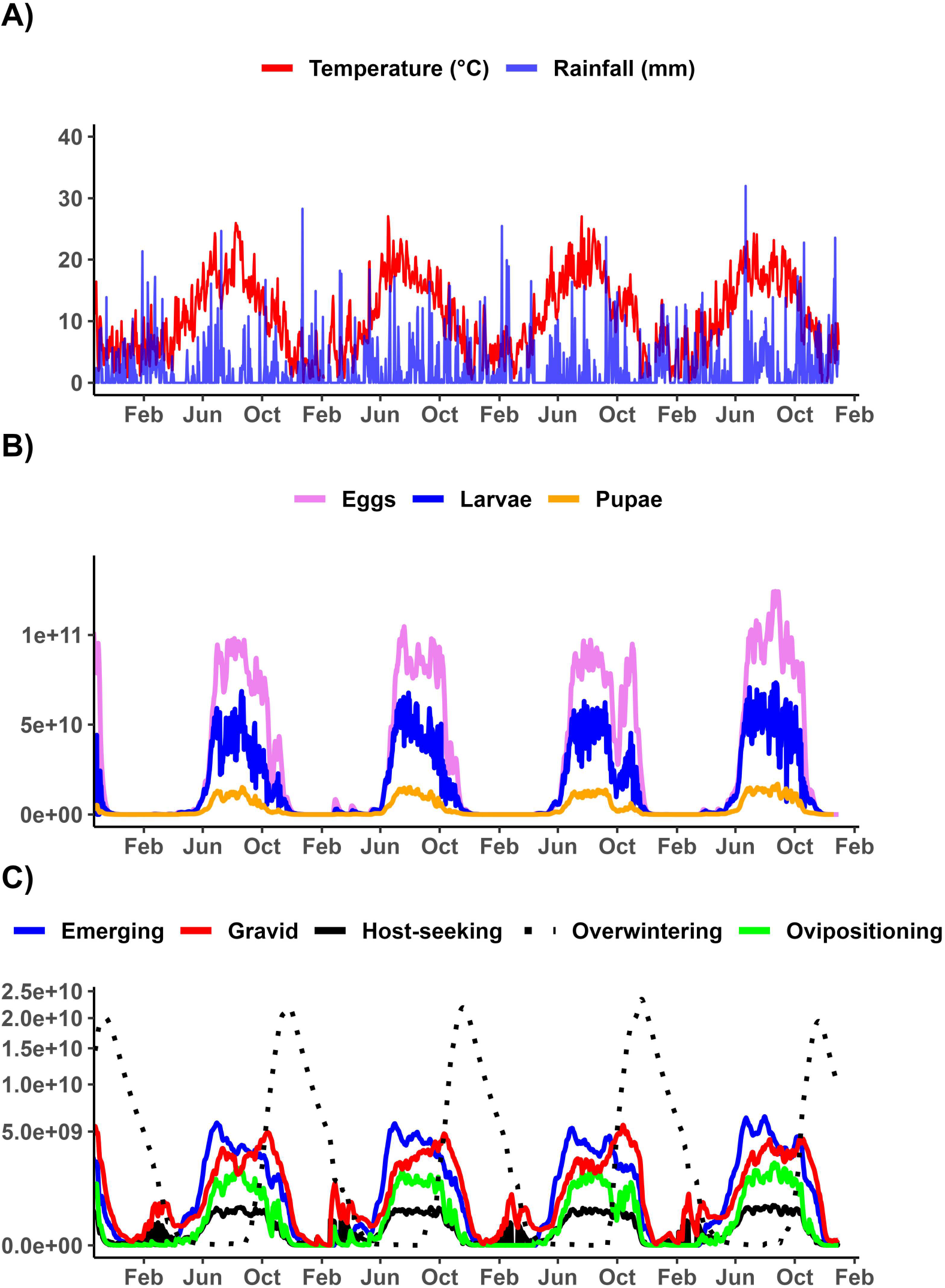
Model output for a single sampling site in Hamburg, Germany, between the years 2020 an 2023, corresponding environmental data A), aquatic life stages (males and females summarised) (B), and adult life stages (nulliparous and parous females summarised) (C).

The timing of the peak in field collected host-seeking females varied by sampling site and year, ranging from the beginning of June to the beginning of October. The average interval between the available field data points was 13 days. The mean offset of abundance peak between the model and field data was 0 days (CI95 = -5.23 - 5.8, median = 0), i.e. on average, the predicted peak day fell within the average interval of field sampling. The predicted and observed start and end of the season also fall within this interval, with a mean offset of 8 days for the start (CI95 = 5.01 - 9.98, median = 0) and -1 day for the end (CI95 = -2.3 - -0.25, median = 0). A significant linear relationship was found between the observed and predicted start of the mosquito biting season (E = 0.75, SE = 0.12, *t* = 6.41, df = 118, *p* < 0.0001, adjusted R^2^ (adj. R^2^) = 0.25; Figure 6). Significant linear relationships were also found for the peak (E = 0.25, SE = 0.11, *t* = 2.18, df = 122, *p* = 0.031, adj. R = 0.03) and for the end of the biting season (E = 0.9, SE = 0.04, *t* = 25.23, df = 119, *p* < 0.0001, adj. R^2^ = 0.84).

**Figure 5:**
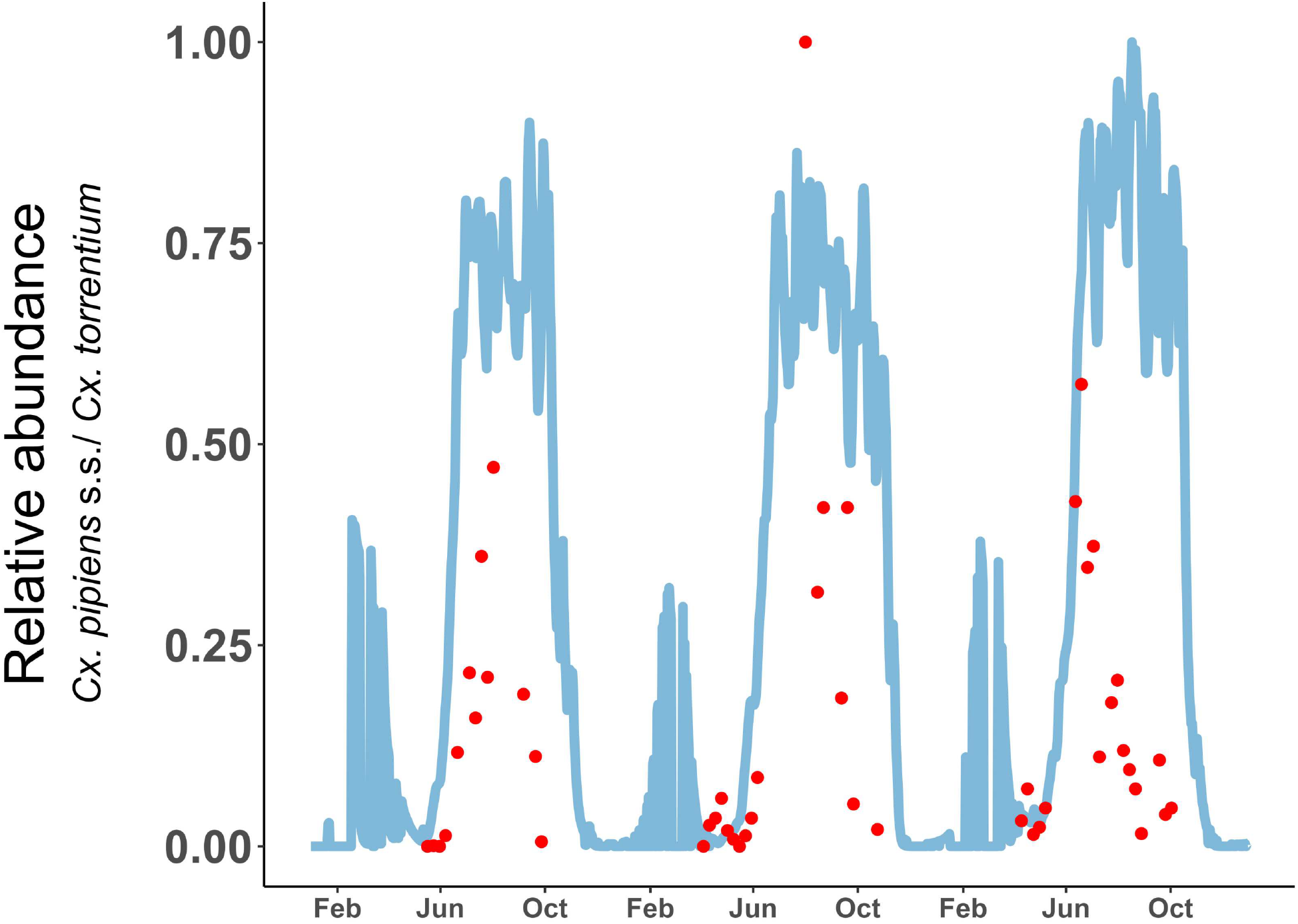
Daily model output of the number of host-seeking *Cx. pipiens* s.s./*Cx. torrentium* (blue) and weekly averaged field data (red) for a single sampling site in Hamburg, Germany between year 2021 and 2023.

**Figure 6:**
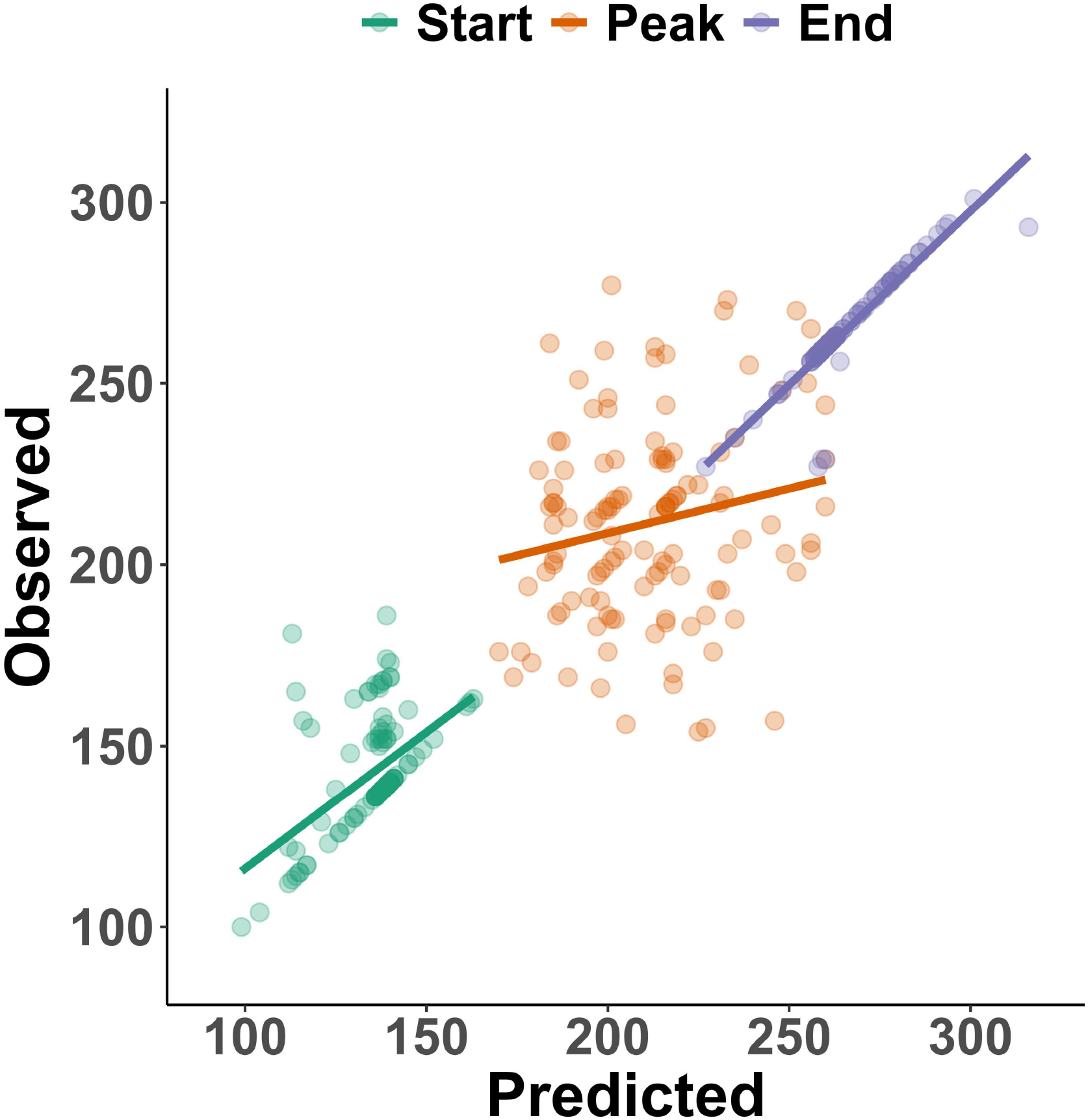
Scatterplot showing the field observed and model predicted start, peak and end of the mosquito biting season including the corresponding linear regressions.

### 3.2. Consecutive generations per season

The calculation of the theoretical maximal number of consecutive generations per year for *Cx. pipiens* s.s./*Cx. torrentium* resulted in a range of 1 to 10 generations per year over Europe, with an average of 7 generations (Figure 7). For example, for Spain, 6 to 10 generations were possible, with an average of 9. In Sweden, 3 to 8 generations per year were calculated with an average of 6 and the number of generations in Germany ranged from 5 to 9, with an average of 8.

**Figure 7:**
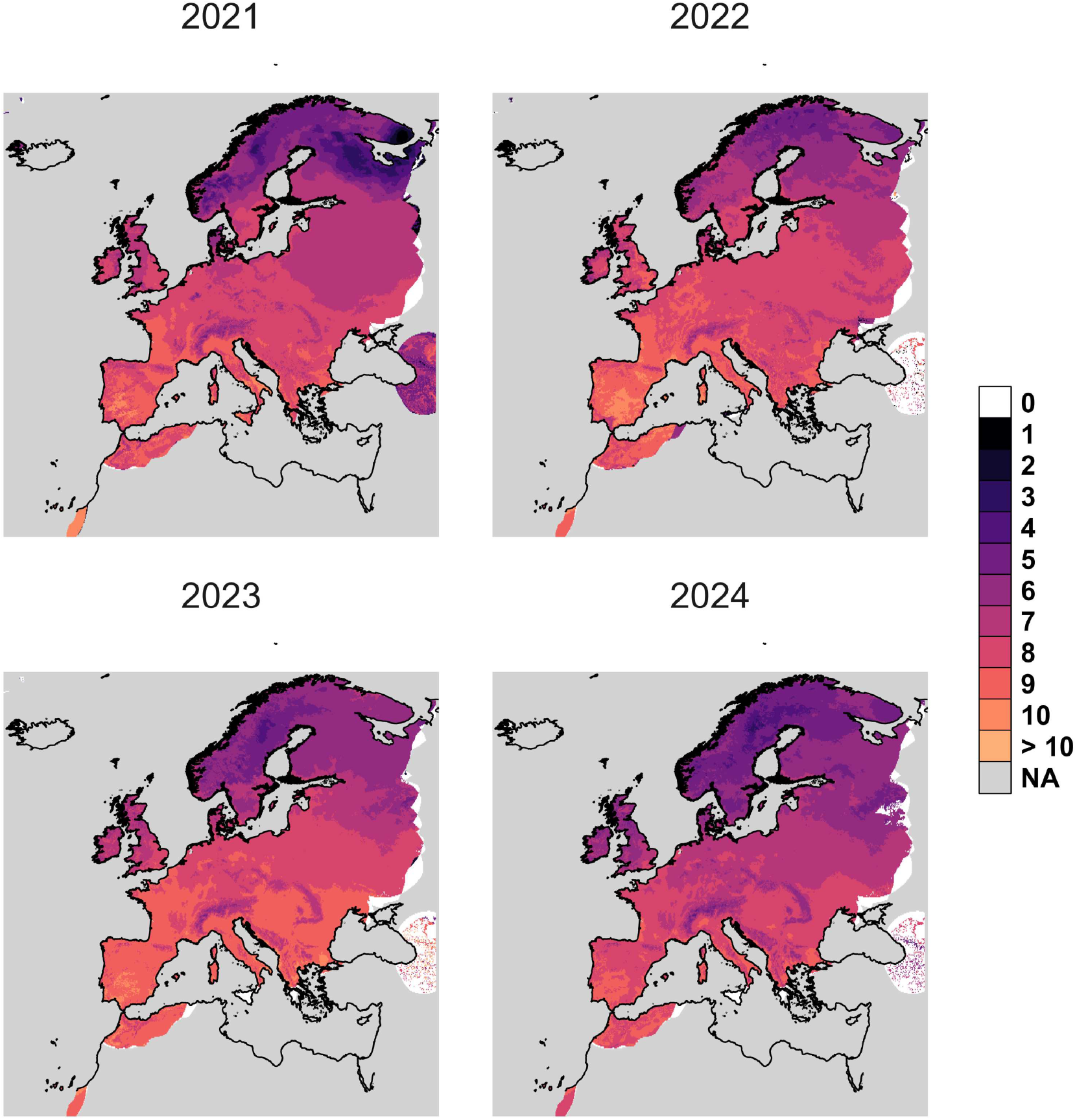
Raster map of maximum number of consecutive generations per year for *Cx. pipiens* s.s./*Cx. torrentium* over Europe

### 3.3. Sensitivity analysis

In the sensitivity analysis, we observe that especially the egg, larval and pupal development rate such as the host-seeking, diapause entering rate and the number of females each male inseminates are positively correlated to the peak of the population of host-seeking adults (PRCCs > 0.2), indicating a strong contribution to the timing of the peak of host-seeking adults (Figure 8). A strong negative impact (PRCCs < -0.2) on the abundance peak was found for the larval and pupal mortality rate and the mortality of adult females. The egg development rate has the highest contribution to the peak (PRCCs = 0.51), followed by the larval development rate with 0.50. The strongest negative impact on the peak in abundance was shown in the mortality rate of adult female mosquitoes (PRCCs = -0.74). We did not observe any strong differences between the sensitivity analysis regarding the peak in abundance compared to the overall phenology (Supplementary table S2).

**Figure 8:**
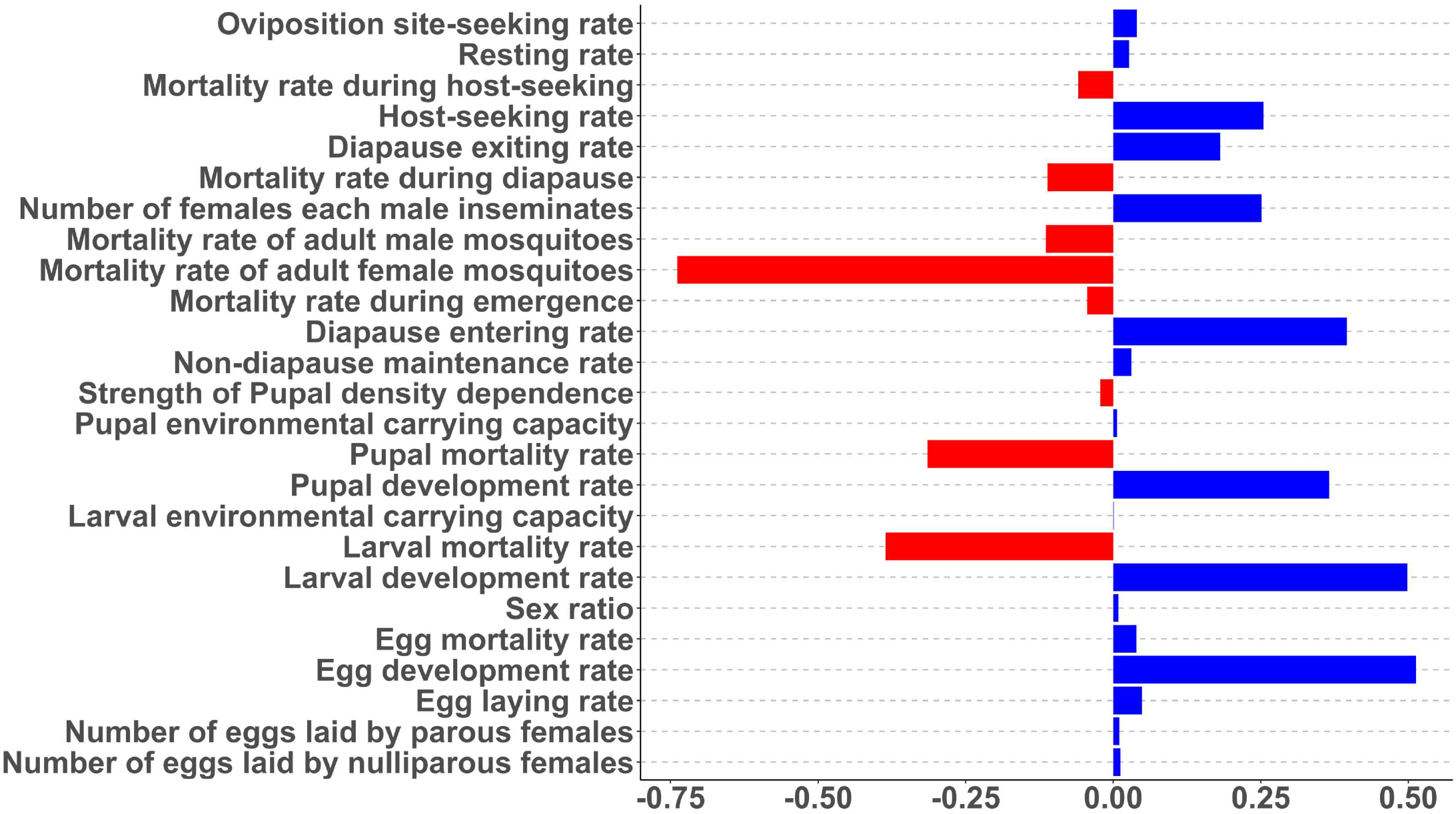
Sensitivity analysis for the peak of host-seeking females

**Figure 9:**
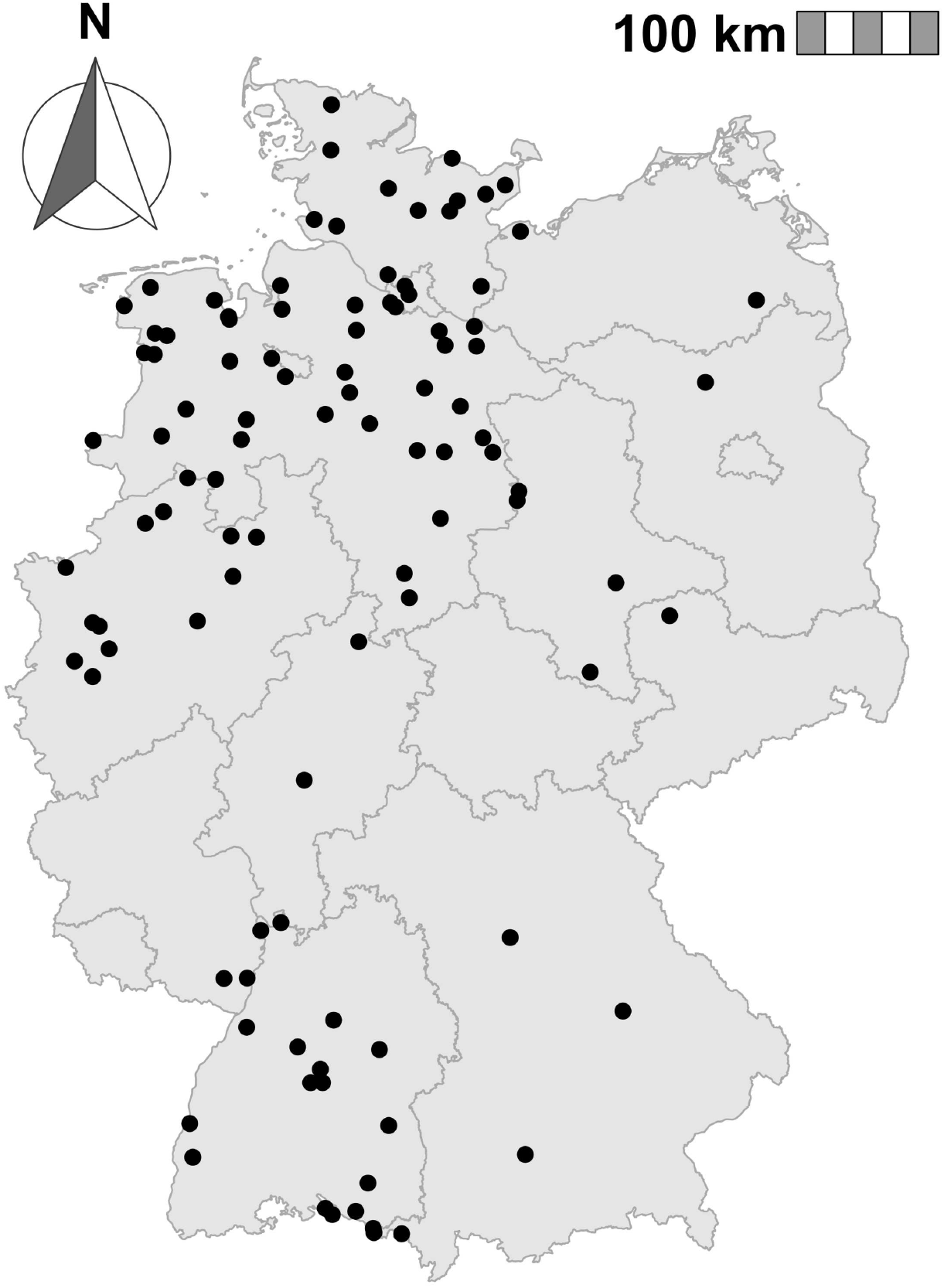
Map of Germany containing all sampling sites used to validate the process-based model

## 4. Discussion

Our updated model with adapted temperature-dependent mortality and transition rates and overwintering compartment was able to predict the seasonal dynamics of *Cx. pipiens* s.s./*Cx. torrentium* for a temperate region. Although we observed strong site-specific differences in mosquito phenology the developed process-based model was able to capture a huge proportion of the observed variability. The newly added compartments, to integrate the overwintering period of the mosquitoes, successfully captured the mosquitoes’ behaviours observed in the field. Female *Cx. pipiens* s.s./*Cx. torrentium* overwinter, while the males do not survive (Sauer et al., 2022, 2023). The inseminated females overwinter in burrows, barns and cellars and are the ones that start the new population by laying eggs in the beginning of the favourable season, after successfully feeding on a host. The photoperiod is considered to be the main trigger for the induction of the diapause (Spielman and Wong, 1973; Field et al., 2022). However, only during low temperatures, the threshold of less than 15 h daytime was found to induce diapause under laboratory conditions. Previous publications either relied on predefined favourable and unfavourable seasons to simulate over-wintering behaviour (Ezanno et al., 2015), or, when diapause was explicitly modeled, it was triggered during a nulliparous stage rather than the larval stage (Andrade et al., 2025). Therefore, to the authors knowledge, this is the first biologically accurate process-based modelling approach to overwintering *Culex* mosquitoes. However, there are important knowledge gaps to fully capture the overwintering of *Culex* mosquitoes. This applies in particular to the termination of diapause at the end of the unfavourable season, where more research is needed to fully grasp the relationship of temperature and photoperiod, terminating diapause (Onyeka and Boreham, 1987; Nelms et al., 2013). The start point of the season can be of great importance for vector control measures, since larvicide treatments during the beginning of the favourable season can have lasting impact on the mosquito population (Runge et al., 2021). Previous published models either estimated a too early start of the season compared to field data (Marini et al., 2016), or overcame the issue by predefining favourable and unfavourable seasons (Tran et al., 2013; Ezanno et al., 2015). In contrast, these studies depicted the end point of the season better than the start point (Marini et al., 2016). Our model was able to predict the start, end and peak of the *Cx. pipiens* s.s./*Cx. torrentium* population in Germany. The inclusion of the overwintering stage allowed for high precision in predicting the start and end of the season. Since photoperiod dynamics remain constant from year to year, only temperature thresholds can postpone the induction or termination of diapause, thereby shortening or extending favourable and unfavourable seasons. The temperature-dependent transition rates are designed to produce biologically meaningful values across the entire viable temperature range, with sigmoid functions reaching an upper plateau around the temperature optima. Previously used transition rates only delivered meaningful rates in a temperature range suitable to the respective study area. On this basis calculated maximum number of consecutive mosquito generations has the advantage that it does not only consider the beginning and ending of the favourable season but incorporates meteorological data of the entire season. Previous studies also highlighted the importance of the seasonal length regarding disease transmission risk (Ewing et al., 2016). A longer biting season is associated with a higher transmission risk. Furthermore, the vertical transmission of a pathogen by an infected mosquito to its offspring, plays a role in the persistence of WNV in temperate regions (Anderson and Main, 2006). Therefore, this process-based model and the extracted maximum number of consecutive generations gives a framework for a more realistic assessment of the local virus transmission risk.

Although not all sampling sites showed a significant linear relationship, the LMM over all sampling sites demonstrated a significant relationship between the number of predicted and actually captured host-seeking females. At the same time, we have shown that the mosquito phenology is highly temperature-dependent, although temperature alone explains less variation in the data set compared to the process-based model. The ability to reliably predict the population peak is an important feature of our model, e.g. to inform pest control management about the best time point for pesticide application. While the linear relationship between the observed and predicted start and end of the mosquito biting season was highly significant, the relationship of the peaks yet being significant showed little explanatory power. Nevertheless, when considering the average interval between observations of 13 days, 38.71% of all sampling sites fall within the interval. The phenology described by the process-based model shows an increase in abundance followed by a plateau caused by the larval and pupal density dependence. It should be noted that mosquito data collected from the field can be easily biased by various factors such as sampling rhythms, weather conditions, trap position or errors by the volunteers running the trap. This can lead to additional challenges in the validation of process-based models. Previous publications addressed such issues by averaging the mosquito collection data over several traps sampled on a weekly or biweekly basis, used for validation on a regional scale (Tran et al., 2013; Ezanno et al., 2015; Marini et al., 2016). In our study, we validated the model based on a large-scale data set from 116 sampling sites across Germany. Thereby, we did not average the data from different sampling sites to preserve the full variability of the mosquito collection data, allowing a detailed assessment of the model accuracy.

A process-based mosquito population model can be a very useful tool as it can provide valuable ecological insights, e.g. start, end and peak of the population or the potential maximum number of consecutive generations per season. Several parameters need further investigation to improve the accuracy of our process-based model. On the one hand, factors such as host-seeking rate or environmental carrying capacities rely on expert estimation, in contrast to temperature-dependent transition rates and mortality rates during the aquatic life stages, which could be at least experimentally measured in climate chambers. Additional compartments for process-based mosquito population models could be considered, i.e. life stages such as resting adult male mosquitoes, however, we still lack the necessary ecological knowledge. Furthermore, the model does not differentiate between *Cx. pipiens* s.s. and *Cx. torrentium*, two different species which probably have different temperature-dependencies (Zittra et al., 2016). However, especially the knowledge of the thermal biology of *Cx. torrentium* is scarce and it is not yet possible to separate these in this model approach. In addition, field data are missing for validation. It has been shown that temperature is the most important climatic factor in driving the phenology of mosquitoes (Lee et al., 2022), but other factors such as photoperiod, wind speed or direction, humidity, and rainfall also influence their general activity (Spielman and Wong, 1973; Camargo et al., 2021).

items, please use

## Supporting information

Linear relationship for each sampling site including the number of sampling days and overall captured specimen

Sensitivity analysis

## 5. Acknowledgements

We gratefully acknowledge the voluntary helpers who supported the field work, and Lisa J. Winter for her help during laboratory work. This work was supported by the Federal Ministry for the Environment, Nature Conservation, Nuclear Safety and Consumer Protection (Grant Number 3721484020), the Federal Ministry of Education and Research (Grant Number 01Kl2022), and the German Research Foundation (Grant Number JO 1276/5-1).

## 5. Availability of data and materials

Validation data and a minimal reproducible example are available under https://github.com/Silence1490/A-process-based-model-simulating-the-life-cycle-of-Culex-pipiens-s.s.-Cx.-torrentium-in-Germany

## 7. Authorship contribution

Conceptualization: LR, PD, FGS, RL; data collection: LR, PD, SMMA, HJ, TS,FGS, LR; data analysis: LR, PD, FGS, RL; first draft: LR, FGS, RL; writing and editing: all authors.

## Notes

https://github.com/Silence1490/A-process-based-model-simulating-the-life-cycle-of-Culex-pipiens-s.s.-Cx.-torrentium-in-Germany

### Competing Interest Statement

The authors have declared no competing interest.

### Summary of Updates

Correction of minor errors, inclusion of Overwintering behaviour and changes in visualisation.

